# A fear conditioned cue orchestrates a suite of behaviors

**DOI:** 10.1101/2022.08.05.502178

**Authors:** Amanda Chu, Christa B. Michel, Nicholas T. Gordon, Katherine E. Hanrahan, Aleah M. DuBois, David C. Williams, Michael A. McDannald

## Abstract

Pavlovian fear conditioning has been extensively used to study the behavioral and neural basis of defensive systems. In a typical procedure, a cue is paired with foot shock, and subsequent cue presentation elicits freezing, a behavior theoretically linked to predator detection. Studies have since shown a fear conditioned cue can elicit locomotion, a behavior that - in addition to jumping, and rearing - is theoretically linked to imminent or occurring predation. A criticism of studies observing fear conditioned cue-elicited locomotion is that responding is non-associative. We gave 24 rats (12 female) Pavlovian fear discrimination over a baseline of reward seeking. The within-subjects procedure had full controls for associative learning, consisting of three cues predicting unique foot shock probabilities: danger (*p*=1), uncertainty (*p*=0.25), and safety (*p*=0). TTL-triggered cameras captured 5 behavior frames/s prior to and during cue presentation. We scored 86,400 frames for nine discrete behaviors spanning reward, passive fear, and active fear. Temporal ethograms show that a fear conditioned cue elicits locomotion, jumping, and rearing that is maximal towards cue offset, when foot shock is imminent. A fear conditioned cue further suppresses reward-related behavior, and elicits freezing in a sex-specific manner. The differing temporal profiles and independent expression of these behaviors reveal a fear conditioned cue to orchestrate a rich and intricate suite of behaviors.

## Introduction

Animals evolved defensive systems to detect and avoid predation. The predatory imminence continuum (PIC), a prominent theory of defensive behavior, identifies three defensive modes based on the proximity to predation: pre-encounter (leaving the safety of the nest), post-encounter (predator detected), and circa-strike (predation imminent or occurring) (Fanselow and Lester, 1988). Pavlovian fear conditioning has been extensively used to reveal the behavioral and neural underpinnings of defensive systems in rats (Bolles and Collier, 1976; Fanselow, 1993; Killcross et al., 1997; McNally et al., 2011). In a typical Pavlovian fear conditioning procedure, a rat is placed in a neutral context and played an auditory cue whose termination coincides with foot shock delivery. Each PIC mode is characterized by a unique set of behaviors and, critically, each mode is thought to be captured by a unique epoch of a Pavlovian fear conditioning trial (Fanselow et al., 2019). The post-encounter mode is characterized by freezing, and is captured by cue presentation. Circa-strike is characterized by locomotion, jumping, and rearing, and is captured by shock delivery.

Freezing to a fear conditioned cue may be the most ubiquitous finding in all of behavioral neuroscience (Blanchard and Blanchard, 1969; Bolles and Collier, 1976; Maren et al., 1997; Anagnostaras et al., 1999; Wilensky et al., 1999; Quirk, 2002; Koo et al., 2004; Rogers and Kesner, 2004; Iordanova et al., 2006; Shumake et al., 2014; Foilb et al., 2016; Furlong et al., 2016). The relationship between freezing and Pavlovian fear conditioning is so strong that failing to observe freezing in defensive settings has been used to support assertions that Pavlovian fear conditioning did not occur (Zambetti et al., 2021). Cued fear as freezing has been further entrenched by historical observations that locomotion, jumping, and rearing (theorized circa-strike behaviors) are not elicited by fear conditioned cues (Fanselow et al., 2019). Instead, active defensive behaviors are restricted to shock delivery (Fanselow, 1982) or to other sudden changes in stimuli (Fadok et al., 2017; Totty et al., 2021). Yet, locomotion, jumping, and rearing all readily occur in defensive settings (Blanchard et al., 1986; Dielenberg and McGregor, 2001) Most relevant, a fear conditioned cue can elicit locomotion, rapid forward movements termed ‘darting’ (Gruene et al., 2015; Mitchell et al., 20_22_).

The ability of a fear conditioned cue to elicit locomotion has been called into question (Trott et al., 20_22_). Trott et al. noted that in prior studies locomotion was greatest at cue onset – the time point most distal from shock delivery (Gruene et al., 2015; Fadok et al., 2017). Moreover, prior studies did not use associative controls (but see Totty et al., 2021) – essential to making claims that cue-elicited behaviors were due to a predictive relationship with foot shock. Using between-subjects designs in mice, Trott et al. ascribe the majority of cue-elicited locomotion to non-associative cue properties. The foundational study demonstrating the need for proper associative controls in *any* form of conditioning used Pavlovian fear conditioning (Rescorla, 1967). Not just all-or-none, the magnitude of a fear conditioned, cue-elicited response can scale with foot shock probability (Rescorla, 1968; Ray et al., 2020). Rescorla 1968, and many foundational associative learning studies (Kamin, 1969; Rescorla and Wagner, 1972), relied on experiments that did not measure ‘fear’ with freezing, but with suppression of operant responding for reward (now termed conditioned suppression) (Estes and Skinner, 1941). Drawing from Rescorla 1968, our laboratory has devised a robust, within-subjects Pavlovian fear conditioning procedure in which three cues predict unique foot shock probabilities: danger (*p*=1), uncertainty (*p*=0.25), and safety (*p*=0). Measuring conditioned suppression we consistently observe complete behavioral discrimination: danger elicits greater suppression than safety, and uncertainty elicits suppression intermediate to danger and safety (Wright et al., 2015; DiLeo et al., 2016; Walker et al., 2018; Ray et al., 2022).

The goal of the current experiment was to construct comprehensive, temporal ethograms of rat behavior during Pavlovian fear conditioning. This would allow us to determine what behaviors come under the control of a fear conditioned cue, and how these behaviors are temporally organized. We had the ability to reveal freezing as the exclusive conditioned behavior, as prior studies have found positive relationships between conditioned freezing and conditioned suppression (Bouton and Bolles, 1980; Mast et al., 1982). Yet, we also had the ability to detect additional behaviors, as brain manipulations that impair conditioned freezing can have little or no impact on conditioned suppression (McDannald, 2010; McDannald and Galarce, 2011). A subgoal was to compare behaviors elicited by the deterministic danger cue, and the probabilistic uncertainty cue.

Twenty-four rats (12 female) received Pavlovian fear discrimination. TTL-triggered GigE cameras were installed in behavioral boxes and programmed to capture frames at subsecond temporal resolution prior to and during cue presentation. 86,400 frames were hand scored for nine discrete behaviors reflecting reward (Holland, 1977), passive fear (Blanchard and Blanchard, 1969; Fanselow, 1982), and active fear (Blanchard et al., 1986; Dielenberg and McGregor, 2001; Gruene et al., 2015). Complete temporal ethograms were constructed for danger, uncertainty, and safety during early, middle and late conditioning sessions. The associative nature of behavioral responding was determined by comparing the danger cue to baseline and safety – the non-associative control. The temporal profile of responding was determined by tracking behavior change over cue presentation.

## Results

### Conditioned suppression reveals complete discrimination

Twenty-four Long Evans rats (12 females) were trained to nose poke in a central port for food reward. Nose poking was reinforced on a 60-s variable interval schedule throughout behavioral testing. Independent of the poke-food contingency, auditory cues were played through overhead speakers, and foot shock delivered through the grid floor (Figure 1A). The experimental design consisted of three cues predicting unique foot shock probabilities: danger (*p*=1), uncertainty (*p*=0.25), and safety (*p*=0) (Figure 1B). Behavior chambers were equipped with TTL-triggered cameras capturing 5 frames/s starting 5 s prior to cue presentation and continuing throughout the 10-s cue. TTL-triggered capture yielded 75 frames per trial, and 1200 frames per session. We aimed to capture 28,800 frames each session (1200 frames x 24 rats).

**Figure 1.**
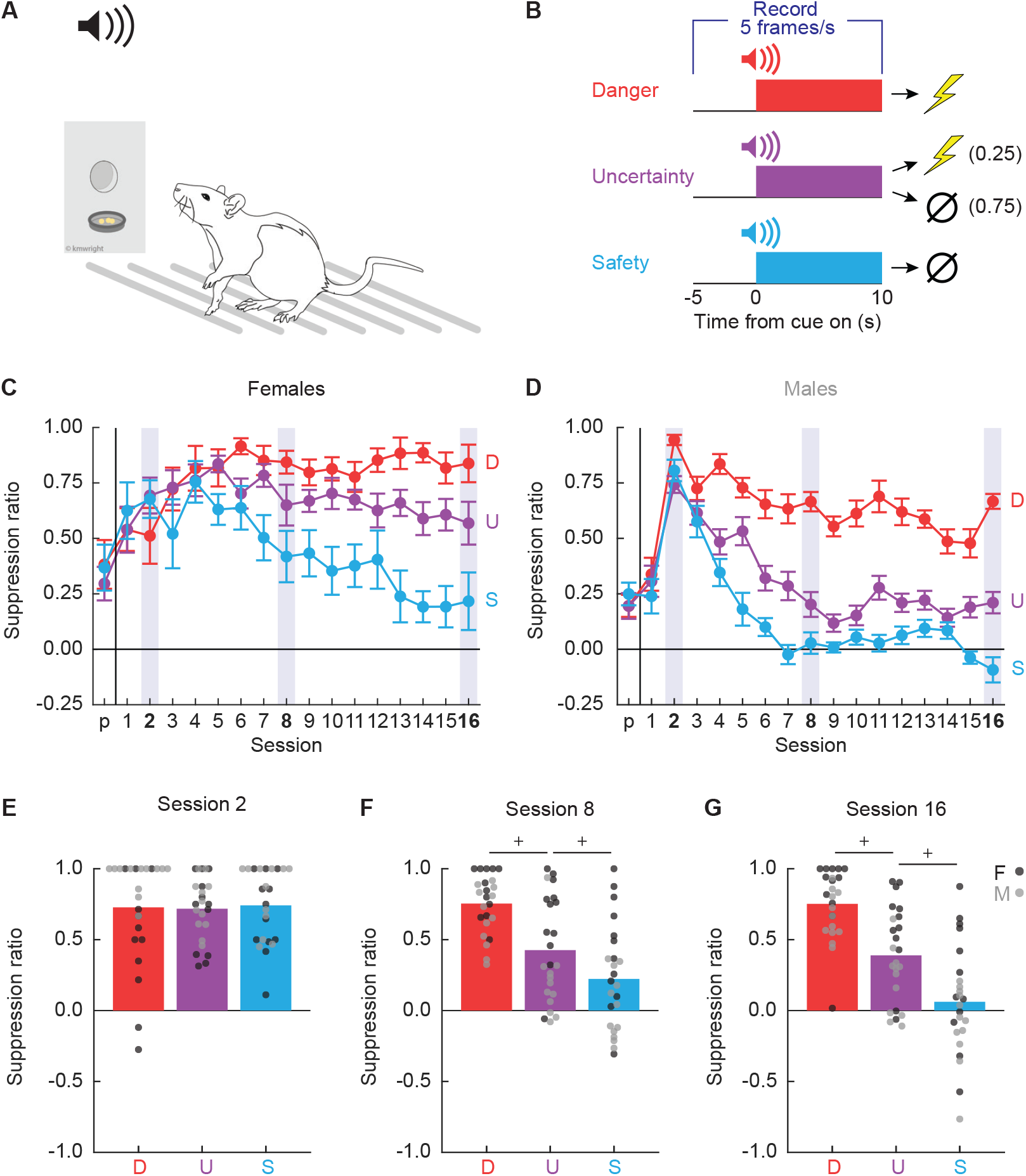
Experimental design and nose poke suppression. **(A)** Conditioned suppression procedure during which rats nose poke for food, while cues are played overhead and shocks delivered through floor. **(B)** Fear discrimination consisted of 10-s auditory cues predicting unique foot shock probabilities: danger (red; *p*=1), uncertainty (purple; *p*=0.25), safety (blue; *p*=0). Five video frames were captured per second, starting 5-s prior to cue onset and continuing through cue presentation. Mean ± SEM suppression ratios for danger (red), uncertainty (purple), and safety (blue) from pre-exposure through discrimination session 16 are shown for **(C)** females, and **(D)** males. Mean + individual suppression ratios for each cue are shown for **(E)** session 2, **(F)** session 8, and **(G)** session 16. Individuals represented by black (female) and gray (male) dots. ^+^95% bootstrap confidence interval does not contain zero.

Our laboratory routinely observes complete behavioral discrimination between danger, uncertainty, and safety in female and male rats measuring conditioned suppression (Walker et al., 2018; Wright et al., 2019; Ray et al., 20_22_). Suppression ratios are calculated using baseline and cue nose poke rates: (baseline - cue) / (baseline + cue). Suppression ratios provide a continuous behavior measure, from no suppression (ratio = 0) to total suppression (ratio = 1). To determine if we observed complete behavioral discrimination in these 24 rats, we performed ANOVA for suppression ratios [factors: cue (danger vs. uncertainty vs. safety), session (17 total: 1 pre-exposure and 16 discrimination), and sex (female vs. male)]. Complete behavioral discrimination emerged over testing (Figure 1C, D). ANOVA found a significant main effect of cue and a significant cue x session interaction (Fs > 6, *p*s < 0.0001; see Table 1 for specific values). Sex effects were apparent; ANOVA found a significant main effect of sex, as well as a significant cue x sex interaction and a cue x session x sex interaction (Fs > 3, *p*s < 0.05; Table 1). Female suppression ratios were higher to each cue across all discrimination sessions: danger (t_22_ = 3.36, *p* = 0.003), uncertainty (t_22_ = 7.14, *p* = 3.67 × 10^−7^), and safety (t_22_ = 4.40, *p* = 0.0002).

**Table 1.**
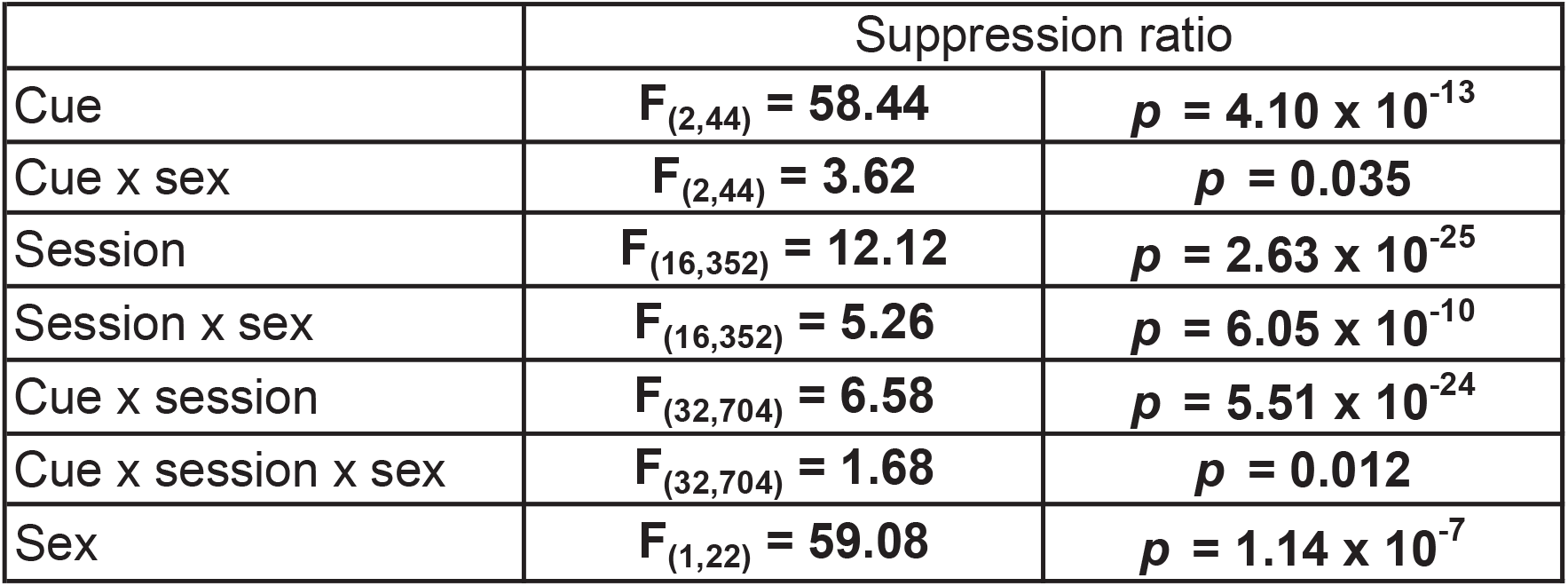
ANOVA results for suppression ratio. Significant main effects and interactions are bolded.

|  | Suppression ratio |  |
| --- | --- | --- |
| Cue | $F_{(2,44)} = 58.44$ | $p = 4.10 \times 10^{-13}$ |
| Cue x sex | $F_{(2,44)} = 3.62$ | $p = 0.035$ |
| Session | $F_{(16,352)} = 12.12$ | $p = 2.63 \times 10^{-25}$ |
| Session x sex | $F_{(16,352)} = 5.26$ | $p = 6.05 \times 10^{-10}$ |
| Cue x session | $F_{(32,704)} = 6.58$ | $p = 5.51 \times 10^{-24}$ |
| Cue x session x sex | $F_{(32,704)} = 1.68$ | $p = 0.012$ |
| Sex | $F_{(1,22)} = 59.08$ | $p = 1.14 \times 10^{-7}$ |

Sex differences in body weight and baseline nose poke rate existed prior to and throughout discrimination, with males weighing more and poking more than females (Figure 1 – Supplemental Figure 1). It is therefore possible that sex indirectly moderates conditioned suppression through effects on body weight or baseline nose poke rate. To determine this, we performed analysis of covariance (ANCOVA) for suppression ratios [factors: cue (danger vs. uncertainty vs. safety) and session (17 total: 1 pre-exposure and 16 discrimination)] using body weight or baseline nose poke rate as the covariate. ANCOVA with body weight found neither a significant body weight x cue interaction (F_(2,44)_ = 2.97, *p*=0.062) nor a significant body weight x cue x session interaction (F_(32,704)_ = 1.40, *p*=0.074). However, ANCOVA with baseline nose poke rate found a significant baseline x cue interaction (F_(2,44)_ = 5.49, *p*=0.007) but not a significant baseline x cue x session interaction (F_(32,704)_ = 0.79, *p*=0.79). Irrespective of sex, higher baseline nose poke rates predicted greater discrimination of danger and uncertainty (Figure 1 - Supplemental Figure 2).

Constructing behavioral ethograms for all 16 discrimination sessions would have required hand scoring 460,800 frames. To make scoring feasible and capture the emergence of discrimination, we selected sessions 2, 8, and 16. Suppression generalized to all cues during Session 2 (Figure 1E). Behavioral discrimination emerged by session 8 (Figure 1F), and was at its most complete during session 16 (Figure 1G). Patterns were confirmed with 95% bootstrap confidence intervals (BCIs) which found no suppression ratio differences for any cue pair during session 2 (all 95% BCIs contained zero), but differences between all cue pairs during sessions 8 and 16 (no 95% BCIs contained zero).

### Hand scoring with high inter-observer reliability

Frames were hand scored for nine discrete behaviors: cup, freezing, grooming, jumping, locomotion, port, rearing, scaling, and stretching, plus ‘background’ (Definitions in Table 2). Behavior categories and their definitions were based on prior work in appetitive conditioning (Holland, 1977), foot shock conditioning (Fanselow, 1982; Blanchard et al., 1986), as well as our own observations. Representative behavior frames are shown in Figure 2.

**Table 2.**
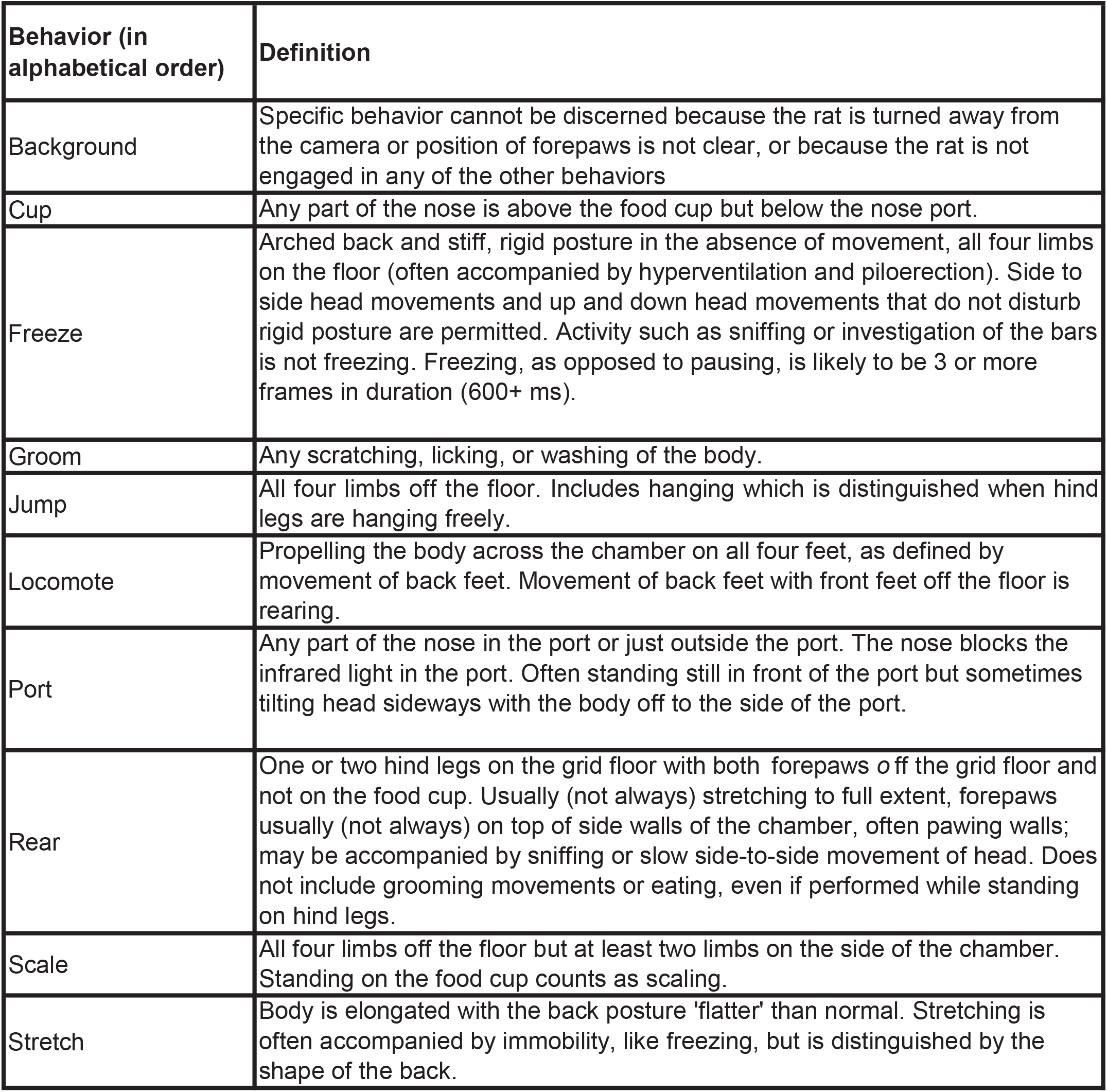
Behavior definitions. Definitions are provided for each behavior scored.

**Figure 2.**
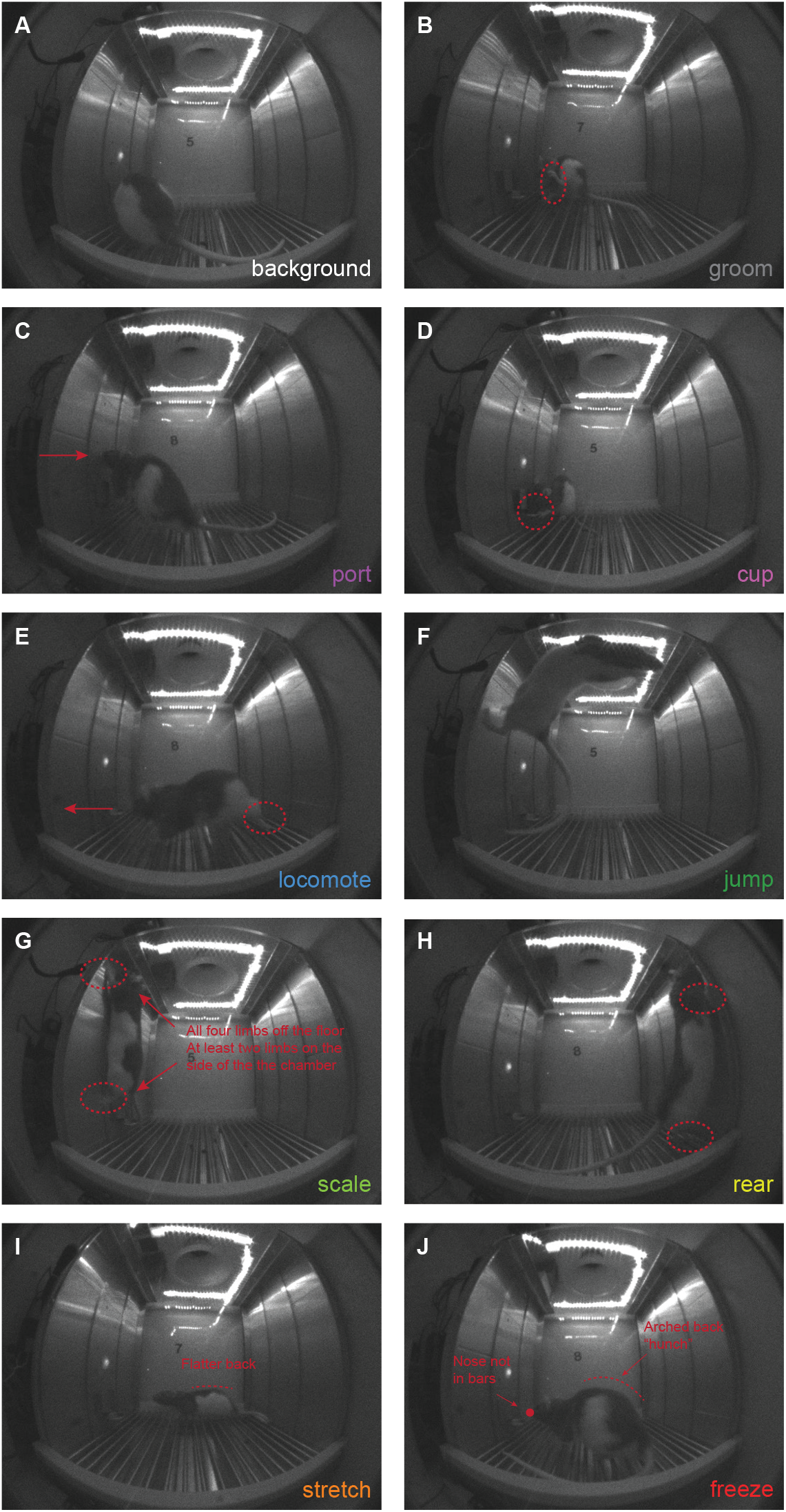
Representative behaviors. Representatives frames are show for: **(A)** background, **(B)** groom, **(C)** port, **(D)** cup, **(E)** locomote, **(F)** jump, **(G)** scale, **(H)** rear, **(I)** stretch, and **(J)** freeze.

Frames were systematically hand scored by five observers blind to rat identity, session number, and trial type (see Materials and Methods for hand scoring approach and trial anonymization). A comparison data set consisting of 12 trials (900 frames) was also scored by each observer. A correlation matrix compared % identical observations for the 900 comparison frames for each observer-observer pair (Figure 3A). Mean % identical observation was 82.83%, with a minimum observer-observer pair agreement of 75.89% and a maximum of 90.56%. Previous studies scoring the presence or absence of freezing have reported inter-observer reliability as an R value: 0.93 (Parnas et al., 2005), 0.96 (Pickens et al., 2010), and 0.97 (Jones and Monfils, 2016). Another study simply reported >95% inter-observer agreement (Badrinarayan et al., 2012). These values exceed our mean % identical observation. However, we hand scored nine discrete behaviors. We observed a negative relationship between the number of behavior categories present and % identical observations (R^2^ = 0.17, *p* = 2.27 × 10^−6^, Figure 3B). Mean percent identical observation was 95% when two behavior categories were present, and 92.5% when three behavior categories were present. Even when eight behavior categories were present, a mean percent identical observation of 78% was achieved. Our approach yielded high inter-observer reliability across trials with few and many behavior categories present. An example of a single-trial ethogram resulting from hand scoring is shown in Figure 3C (female rat, discrimination session 8, uncertainty cue). Videos 1-4 show example danger trials for four different rats (females in Videos 1 & 3, males in Videos 2 & 4).

**Figure 3.**
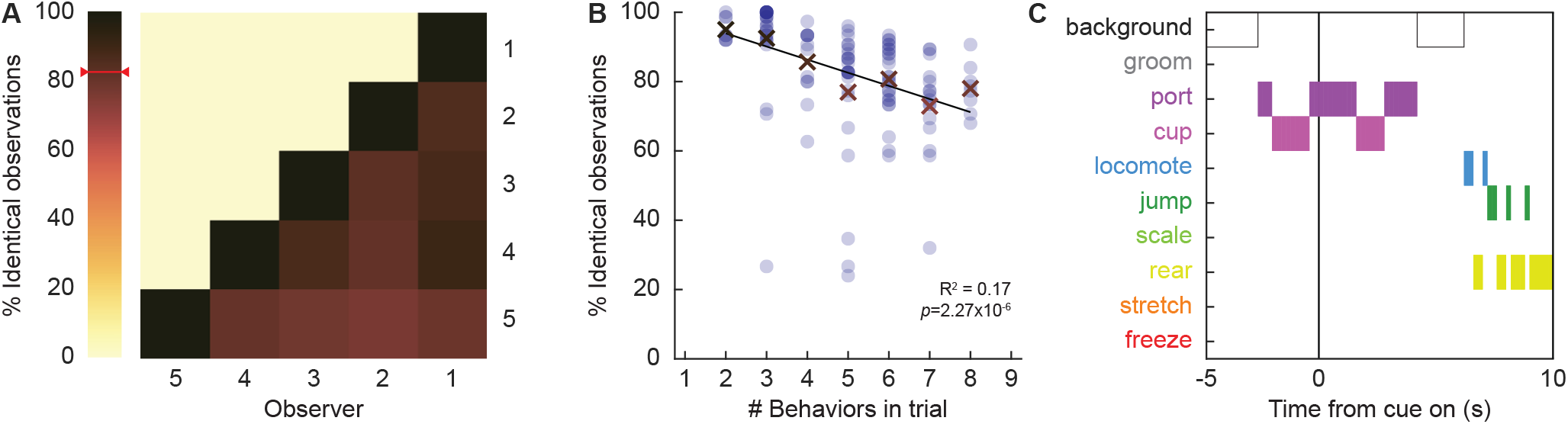
Inter-observer reliability and example ethogram. **(A)** Percentage of identical observations between observer-observer pairs. **(B)** Percentage of identical observations as a function of the number of behaviors present in a trial. **(C)** Example ethogram from a single uncertainty cue presentation, taken from a female during session 8.

### Temporal ethograms reveal shifting behavioral patterns over discrimination

The 86,400 scored frames allowed us to construct temporal ethograms for danger (Figure 4A-C), uncertainty (Figure 4D-F), and safety (Figure 4G-I) during sessions 2 (Figure 4, column 1), 8 (Figure 4, column 2), and 16 (Figure 4, column 3). Shifts in the composition of behavior from baseline to cue presentation were apparent across all ethograms. During session 2 (column 1), behavioral shifts lacked cue-specificity. Temporal ethograms revealed danger, uncertainty, and safety to equally suppress grooming, port, and cup behavior, but increase freezing, and locomotion. Generalized cue control of behavior was supported by multiple analysis of variance (MANOVA) for all nine behavior categories [factors: cue (danger vs. uncertainty vs. safety), time (15 1-s bins: 5-s baseline → 10-s cue), and sex (female vs. male)] revealing a significant main effect of time (F_(126,2772)_ = 2.37, *p*=5.93 × 10^−15^), but neither a significant main effect of cue (F_(18,74)_ = 1.00, *p*=0.47) nor a significant cue x time interaction (F_(252,5544)_ = 1.12, *p*=0.11). Cue-specific shifts in behavior were apparent by session 8 (column 2), and continued to session 16 (column 3). Now, MANOVA revealed significant main effects of cue (session 8, F_(18,74)_ = 3.39, *p*=0.0001; session 16, F_(18,74)_ = 4.44, *p*=0.000002), and significant cue x time interactions (session 8, F_(252,5544)_ = 1.52, *p*=3.31 × 10^−8^; session 16, F_(252,5544)_ = 1.52, *p*=4.74 × 10^−7^).

**Figure 4.**
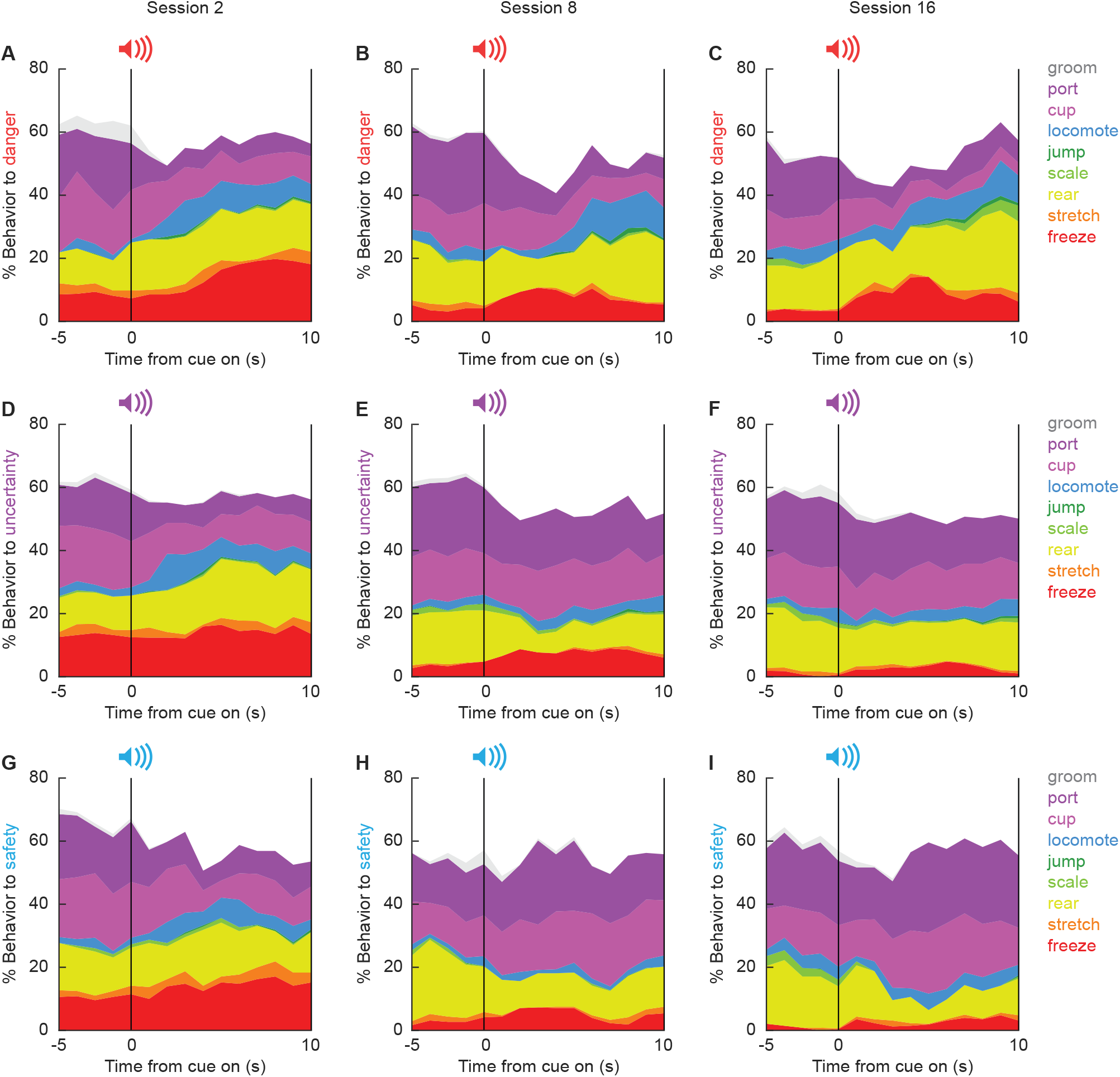
Temporal ethograms. Mean percent behavior from 5s prior through 10-s cue presentation is shown for the danger cue during sessions **(A)** 2, **(B)** 8, **(C)** and 16; the uncertainty cue during sessions **(D)** 2, **(E)** 8, and **(F)** 16; and the safety cue during sessions **(G)** 2, **(H)** 8, and **(I)** 16. Behaviors are groom (gray), port (dark purple), cup (light purple), locomote (blue), jump (dark green), scale (light green), rear (yellow), stretch (orange), and freeze (red).

### Danger orchestrates a suite of behaviors

The central question driving this study is what behaviors come under the specific control of the fear conditioned, danger cue? To determine this, we focused on session 16, when discrimination was at its most complete. We first performed MANOVA for the 5-s baseline period [factors: cue (danger vs. uncertainty vs. safety), time (5, 1-s bins), and sex (female vs. male)]. As expected, MANOVA returned no main effect of cue, time, nor a cue x time interaction (Fs < 1.5, *p*s > 0.1). Univariate ANOVA results were subjected to Bonferroni correction (*p* < 0.0055, 0.05/9 = 0.0055) to account for the nine separately analyses. Like for MANOVA, univariate ANOVA for each of the nine behaviors showed no main effect of cue, time, nor a cue x time interaction. In contrast to all other behaviors, univariate ANOVA for baseline freezing showed a main effect of sex (F(_1,22_) = 10.37, *p*=0.004). ANOVA for freezing across the baseline and cue periods revealed a significant sex x cue x time interaction (F_(28,616)_ = 1.94, *p*=0.003). The unique freezing pattern warrants separate consideration, which we return to later.

MANOVA was then performed for the 10-s cue period [factors: cue (danger vs. uncertainty vs. safety), time (10, 1-s bins), and sex (female vs. male)]. MANOVA returned significant main effects of cue and time, as well as a significant cue x time interaction (Fs >1.3, *p*s < 0.005). Of most interest, univariate ANOVA found a significant main effect of cue for six of the nine behaviors: port (F_(2,44)_ = 32.15, *p*=2.47 × 10^−9^, Figure 5A), cup (F_(2,44)_ = 18.40, *p*=0.00002, Figure 5B), locomote (F_(2,44)_ = 6.33, *p*=0.004, Figure 5C), jump (F_(2,44)_ = 10.90, *p*=0.0001, Figure 5D), rear (F_(2,44)_ = 8.64, *p*=0.001, Figure 5E), and freeze (F_(2,44)_ = 13.86, *p*=0.00002). Danger suppressed port and cup behavior (Figure 5 A, B line graphs), but promoted locomotion, jumping, and rearing (Figure 5 C, D, E line graphs). Danger-specific control of behavior was most apparent in the last 5 s of cue presentation (Figure 5, shaded region).

**Figure 5.**
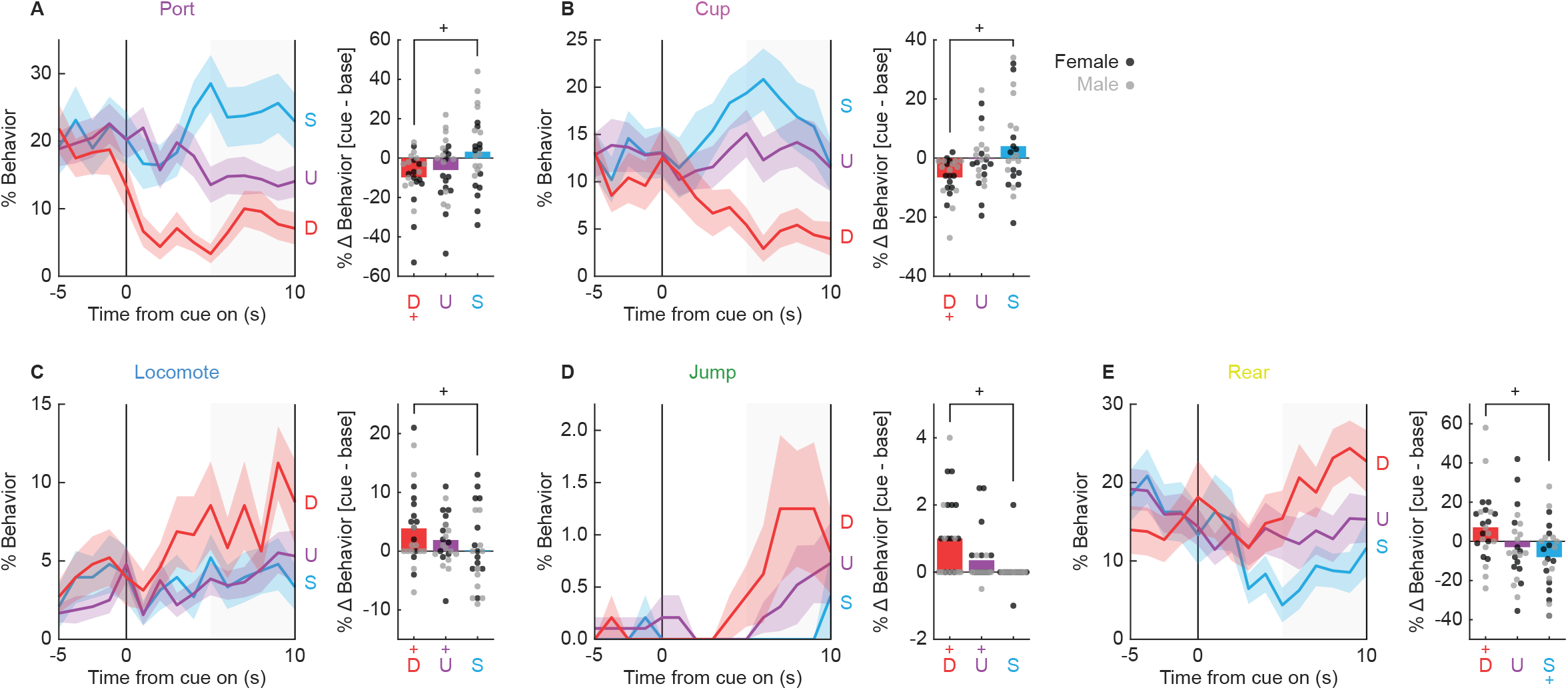
Fear conditioned, cue-elicited behaviors. Line graphs show mean ± SEM percent behavior from 5s prior through 10-s cue presentation for danger (red), uncertainty (purple), and safety (blue) for **(A)** port, **(B)** cup, **(C)** locomote, **(D)** jumping, and **(E)** rearing. Bar plots show mean change in behavior from baseline (5 s prior to cue) compared to last 5 s of cue. Individuals represented by black (female) and gray (male) dots. ^+^95% bootstrap confidence interval for danger vs. safety comparison does not contain zero.

Claiming danger specificity requires that % behavior during the danger cue differs from baseline as well as the safety cue. To test this, we subtracted mean % behavior during the 5-s baseline from mean % behavior during the last 5 s of cue presentation, giving %Δ danger, %Δ uncertainty, and %Δ safety for each subject. We constructed 95% BCIs for each cue/behavior. Confirming danger elicitation of behavior that deviated from baseline, 95% BCIs for %Δ danger did not contain zero for each of the five behaviors (Figure 5). Danger presentation decreased port and cup behavior below baseline, but increased locomotion, jumping, and rearing over baseline. 95% BCIs for %Δ uncertainty revealed increased locomotion and jumping, while 95% BCIs for %Δ safety revealed only decreased rearing. To demonstrate danger-specificity, we subtracted %Δ safety from %Δ danger. We then constructed 95% BCIs for the difference score for each behavior. Confirming danger specificity, 95% BCIs did not contain zero for each of the five behaviors. Thus, danger specifically and selectively suppressed reward-related port and cup behavior, but promoted locomotion, jumping, and rearing.

### Associatively acquired behaviors generalize early

By the end of session 16 each rat had received 96 total foot shocks. It is possible that danger-specific control of multiple behaviors was only observed in session 16 because rats received far more cue-shock pairings than a typical Pavlovian conditioning procedure employs. Session 2 provided a comparison to numbers of cue-shock pairings more typical of fear conditioning studies; rats had received 12 total foot shocks by session’s end. The key question was whether pattern of danger-elicited behaviors in session 2 resembled the pattern in session 16, or if a fundamentally different pattern was observed. To determine this, we performed univariate ANOVA for danger [factors: session (2 vs. 16) and time (15, 1-s bins)] for each of the five behaviors showing session 16 selectivity (Figure 5 – Supplemental Figure 1). Confirming near identical temporal patterns of behavior expression during sessions 2 and 16, ANOVA found no significant session x time interaction for any behavior [port (F(_14,322_) = 0.45, *p*=0.96), cup (F(_14,322_) = 0.61, *p*=0.86), locomote (F(_14,322_) = 1.09, *p*=0.37), jump (F(_14,322_) = 1.23, *p*=0.25), and rear (F(_14,322_) = 0.92, *p*=0.54)]. Thus, danger orchestrated a suite of behaviors even early in discrimination. Recall that early discrimination (session 2) was marked by non-specific cue control of behaviors. This would mean that associatively acquired behaviors initially generalized to uncertainty and safety – and that discrimination consisted of restricting behavior to danger. In support, univariate ANOVA for session 2 [factors: cue (danger vs. uncertainty vs. safety), time (15, 1-s bins), and sex (female vs. male)] found no cue x time interaction for any of the five, danger-specific behaviors (all Fs < 1.2, all *p*s > 0.3).

### Sex informs the temporal pattern of freezing

We return to the case of freezing; the most measured overt fear conditioned behavior. We again focus on session 16 during which discrimination was most complete. Female and male rats differed in the temporal pattern and cue-specificity of freezing. Females showed higher baseline freezing levels, a rapid increase in freezing that was specific to danger in the first 5 s, then became non-specific and declined back to baseline levels in the last 5 s (Figure 6A). By contrast, males show little baseline freezing and danger-specific freezing increases that persisted throughout cue presentation (Figure 6B). Baseline freezing differences were confirmed with independent samples t-test (t_22_ = 3.22, *p*=0.0039; Figure 6C). Confirming sex differences in the temporal pattern of freezing, differential freezing to danger and safety was equivalent in females and males during early cue presentation (t_22_ = 0.02, *p*=0.98; Figure 6D left), but differed during late cue (t_22_ = 2.80, *p*=0.01; Figure 6D right). Generalized freezing to all cues was observed during session 2, with freezing increases more evident in males (Figure 6 – Supplemental Figure 1). Thus, discrimination consisted of restricting freezing to danger in males, and selectively freezing to early danger presentation in females.

**Figure 6.**
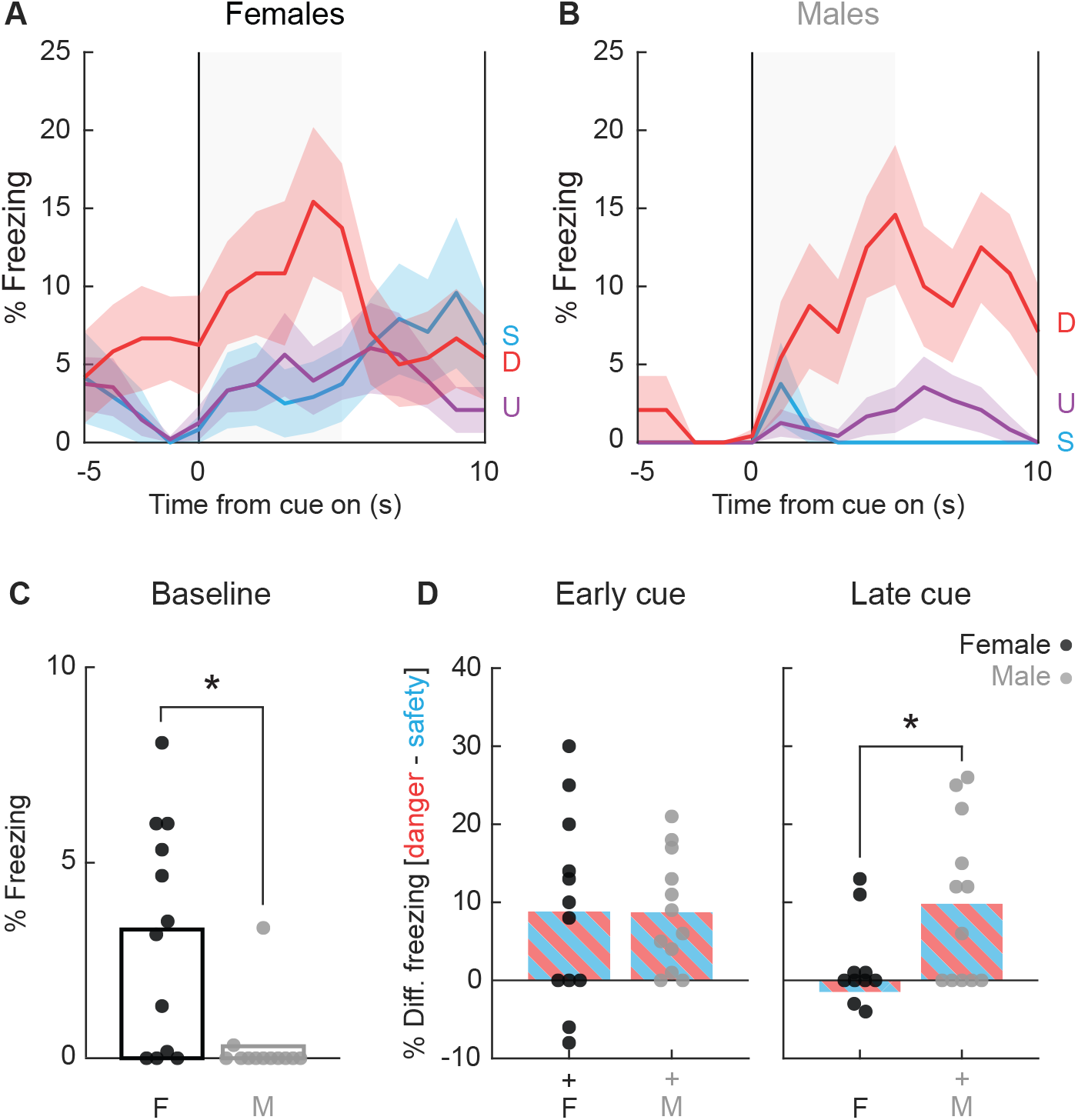
Special case of freezing. Line graphs show mean ± SEM percent freezing from 5s prior through 10-s cue presentation for danger (red), uncertainty (purple), and safety (blue) for **(A)** females and **(B)** males. **(C)** Percent freezing during baseline (5s prior to cue) is shown for females (black) and males (gray). **(D)** Mean differential freezing to danger and safety is shown for females (black, left) and males (gray, right) during early cue (first 5s of cue, left) and late cue (last 5s of cue, right). Mean ± SEM percent freezing change from baseline (5 s prior to cue) compared to last 5 s of danger (red), uncertainty (purple), and safety (blue) for **(E)** females and **(F)** males.

### Danger-elicited behaviors are independently expressed

Danger suppression of reward-related port and cup behaviors could simply be the byproduct of danger-elicited freezing. Such a relationship has previously been reported <u>(Bouton and Bolles,</u> 1980; Mast et al., 1982). To examine the relationship between reward-related behaviors and freezing, in addition to other possible behavior-behavior relationships, we calculated %Δ behavior for early (first 5 s) and late (last 5 s) danger presentation for the six danger-elicited behaviors: cup, port, locomote, jump, rear, and freeze. We constructed 12 × 12 matrices containing the R values (Figure 7A) and *p* values (Figure 7B) for the Pearson’s correlation coefficient for each behavior-behavior comparison during the two danger periods. Surprisingly, a single behavior-behavior relationship was observed during early danger presentation period (Figure 7A, upper left quadrant). Early rearing and early cup behavior were negatively correlated (R = -0.43, *p* = 0.03, but note this would not survive Bonferroni correction). Even more, no behavior-behavior relationships were observed during late danger presentation (Figure 7A, lower right quadrant). These results suggest the six behaviors are more or less expressed independently of one another.

**Figure 7.**
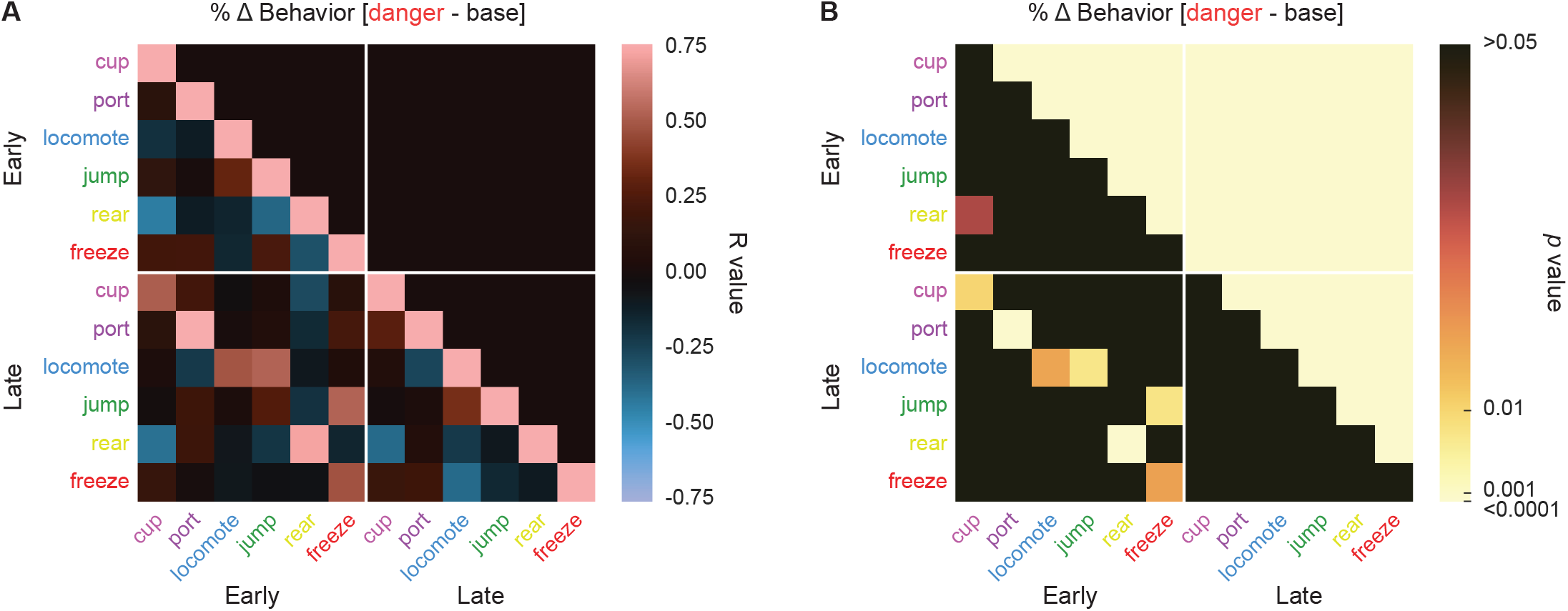
Behavior-behavior correlations. **(A)** A correlation matrix for the six cue specific behaviors (port (dark purple), cup (light purple), locomote (blue), jump (dark green), rear (yellow), and freeze (red) comparing mean percent behavior during early (first 5 s) and late (last 5 s) cue is shown. Lighter red values indicate positive R values, lighter blue values indicate negative R values. Black indicates R = 0. *P* values associated with each associated R value are shown in **(B)**. Black indicates *p* values greater than 0.05, while increasingly lighter values indicate lower *p* values.

Maybe our analysis cannot detect behavior-behavior relationships? To test this, we compared behaviors across the early and late danger periods. Now, the correlation matrix revealed a band of positive R values cutting diagonally across the bottom left quadrant. 5 of the 6 behaviors showed positive early-late relationships with themselves: cup (R = 0.51, *p* = 0.01), port (R = 0.87, *p* = 2.67 × 10^−8^), locomote (R = 0.48, *p* = 0.017), rear (R = 0.71, *p* = 7.92 × 10^−5^), and freeze (R = 0.48, *p* = 0.017) from early to late danger presentation. In other words, changes in cup behavior evident during early danger presentation persisted to late danger presentation. Jumping was an exception to this trend, as there was no relationship between early and late jumping levels to danger. Overall, danger-elicited behaviors were expressed independently of one another.

## Discussion

We set out to quantify behaviors elicited by a fear conditioned, danger cue. Consistent with virtually all studies of Pavlovian fear conditioning (but see Amorapanth et al., 1999), we observed danger-elicited freezing in both female and male rats. Biological sex mediated the temporal expression of freezing, with females showing an inverted ‘V’ pattern in which freezing peaked mid danger cue, while males sustained freezing over the entirety of danger presentation. Yet, freezing was not the dominant danger-elicited behavior. Instead, danger orchestrated a suite of behaviors. Danger suppressed reward behavior directed toward the site of food delivery and the location of the rewarded action, and this suppression was not a byproduct of freezing. Even more, danger elicited ‘active’ behaviors: locomotion, jumping, and rearing.

Before discussing our results further, an important limitation should be raised. 40-50% of frames could not be assigned to a specific behavior and were labeled as background. This was due to three main factors. First, in order to objectively hand score many behaviors, we developed mutually-exclusive, specific definitions. Our strict definitions meant erring on the side of labeling a behavior background if there was uncertainty in judgment. Second, use of a single, side view camera meant the observer could not view a rat’s forelimbs or face when the rat was turned away from the camera. If forelimb and face position could not be determined the frame was labeled background. Finally, transition behaviors (e.g. switching from rearing to locomotion) and other behaviors (e.g. sniffing and turning) that did not fit into one of the nine behavior definitions were labeled background. The upside of this limitation is high confidence in behavior judgments and high inter-rater reliability for those judgments.

Studies assessing auditory fear conditioning in a neutral context routinely report freezing to account for >80% of behavior during cue presentation (Bolles and Collier, 1976; Maren et al., 1997; Anagnostaras et al., 1999; Wilensky et al., 1999; Quirk, 2002; Koo et al., 2004; Rogers and Kesner, 2004; Iordanova et al., 2006; Shumake et al., 2014; Foilb et al., 2016; Furlong et al., 2016). The sheer number of demonstrations, and number of groups independently observing dominant freezing, puts us firmly in the minority. Placing us further in the minority, we observe danger elicitation of ‘active’ behaviors: locomotion, jumping, and rearing. These danger-elicited behaviors are characterized by lateral and vertical movements, polar opposites to the immobility that characterizes freezing.

However, we are not the first group to observe locomotion, jumping, or rearing in defensive settings in rats. Using more traditional Pavlovian fear conditioning designs, Shansky and colleagues have observed ‘darting’, rapid forward movements across the test chamber, to a fear conditioned cue (Gruene et al., 2015). While more readily observed in female rats, darting can be observed in males under certain experimental conditions (Mitchell et al., 20_22_). Our definition of locomotion aligns well with the Shansky lab definition of darting, demonstrating at least three settings in which a fear conditioned cue promotes movement in rats (Totty et al., 2021). Jumping is readily elicited in rats by hypoxia (decreasing oxygen levels in the air) – a life threatening condition (Spiacci et al., 2015). More relevant to our study, the Blanchards readily observed jumping in defensive settings in rats (Blanchard et al., 1986). In their procedure, a rat was placed at the end of an inescapable hallway, then a human experimenter slowly approached. Rats initially froze when the experimenter was distant (1-2 meters away), but switched from freezing to jumping as the experimenter drew near (<1 meter). Our observation of danger-elicited jumping, and its preferential expression at the end of danger presentation, mirrored the defensive jumping pattern observed in the Blanchard’s study.

Rearing may be the least reported behavior in defensive settings. Further, behavioral interpretations of rearing vary widely (C. Lever et al., 2006). Nevertheless, Dielenberg and McGregor found that rats exposed to a recently worn cat collar, with an opposing box to hide in, show ‘vigilant rearing’ to the cat collar (Dielenberg and McGregor, 2001). Rearing was never observed in a control condition. While we cannot claim vigilance, we find that danger promotes rearing. Finally, our temporal ethograms revealed that locomotion, jumping, and rearing were all most prevalent at the end of cue presentation – when foot shock was imminent. This was in contrast to freezing which was prominent during early danger presentation for both females and males, but only shown by males at the end of cue presentation.

Our findings accord well with the PIC theory of defensive behavior (Fanselow and Lester, 1988). PIC theory identifies three defensive modes: pre-encounter (for example, leaving the safety of the nest to forage), post-encounter (predator detected), and circa-strike (predation inevitable or occurring). Analogs to PIC modes are identified in a Pavlovian fear conditioning trial (Fanselow et al., 2019). Pre-encounter mode may correspond to leaving the home cage and being placed in the experimental chamber where foot shocks occur. Post-encounter mode corresponds to presentation of the fear conditioned cue. Circa-strike mode is said to correspond to foot shock delivery. It is argued that circa-strike behaviors (locomotion, jumping, and rearing) are not observed towards the end of danger presentation because rats do not time shock delivery. In support, extending cue duration in traditional cued fear conditioning paradigms does not result in shifts from freezing to locomotion, jumping, and rearing towards cue offset (Fanselow et al., 2019).

Here we find expected patterns of defensive behavior in unexpected epochs of Pavlovian conditioning trials. Early danger freezing by all rats (females and males) gives way to a late mix of danger-elicited behaviors that included locomotion, jumping, and rearing. Why do we observe late danger control of circa-strike behaviors? Hunger and the availability of a rewarded action may provide an impetus for shock timing. Timing would allow rats to minimize the display of defensive behaviors and maximize reward seeking. In support, presenting long duration danger cues in a conditioned suppression settings results in timing of fear responding. With experience, rats show little suppression of reward seeking to danger onset, which ramps towards shock delivery (Rosas and Alonso, 1996). Supporting the minimization of defensive behavior in reward settings, foot shocks signaled by danger will strongly suppress reward seeking only early in fear conditioning. Shock-induced suppression quickly wanes and with experience, shock delivery will paradoxically facilitate reward seeking (Strickland et al., 2021). Shock timing information is readily apparent in the ventrolateral periaqueductal gray, a brain region central to defensive behavior (Fanselow, 1993; Carrive et al., 1997; Mobbs et al., 2007; McDannald, 2010; Tovote et al., 2016; Arico et al., 2017). Populations of ventrolateral periaqueductal gray neurons show little responding upon danger presentation, but ramp firing towards shock delivery (Ozawa et al., 2017; Wright and McDannald, 2019; Wright et al., 2019). Our results support PIC theory of defensive behavior but demonstrate that the relationship between defensive mode and Pavlovian conditioning trial epoch is not fixed, but depends on experimental setting.

A secondary goal was to compare defensive behaviors elicited by a deterministic, danger cue and a probabilistic, uncertainty cue. In our setting, uncertainty is not simply a diminished version of danger. Indeed, uncertainty only promoted a subset of danger-elicited behaviors: locomotion and jumping. Most surprising was the inability of uncertainty to suppress reward behaviors directed towards the food cup and port. This is particularly puzzling because using suppression ratios, we found uncertainty to produce robust suppression of nose poking. What is going on here? Food cup, port and poke behavior lie on a rewarded action continuum. Food cup means the rat is in the area of food delivery – but is most distal from the rewarded action. Port means the rat is around or in the site of the rewarded action, but only poke requires the rat to be fully engaged in performing the rewarded action (nose all the way into the port). Danger suppresses all reward behavior regardless of proximity to rewarded action. By contrast, uncertainty selectively suppresses reward behavior most proximal to the rewarded action.

By comprehensively quantifying behavior and constructing temporal ethograms, we found a fear conditioned cue to independently control at least six distinct behaviors. Though our study was exclusively behavioral, we feel our results have implications for investigations into the neural basis of fear learning and the organization of defensive behavior. Most important is that a fear conditioned cue does not simply elicit freezing. Behaviors elicited by a fear conditioned cue are the product of many factors: species, sex, age, behavioral setting, and experimenter determined parameters (CS/US type, duration, and intensity; trial number, inter-trial interval, and more). In our view, freezing is a common – not dominant – defensive behavior because the field has favored behavioral settings and experimenter determined parameters that maximize the expression of ‘fear’ through freezing. Here we show that a relatively simple modification of the rat’s behavioral setting – access to a rewarded action – is sufficient to de-emphasize freezing and promote the expression of many additional behaviors. Even more, Pavlovian fear conditioning over a baseline of reward seeking reveals a temporally organized sequence of cue-elicited defensive behaviors predicted by PIC theory. The independent expression of these behaviors is appealing for studies attempting to link discrete behavioral sequelae of ‘fear’ to distinct neural circuits; breathing new life into a classic Pavlovian fear conditioning procedure (Estes and Skinner, 1941).

## Figure Titles and Legends

**Video 1**. Behavior during a single danger trial

Video shows the 75 sequential frames for a danger trial. Frames 1-25 are background and 26-75 are danger cue presentation. Observer judgment is shown in the top right for each frame. The specific trial is 23_16_12, (female rat 23, session 16, trial 12).

**Video 2**. Behavior during a single danger trial

Video shows the 75 sequential frames for a danger trial. Frames 1-25 are background and 26-75 are danger cue presentation. Observer judgment is shown in the top right for each frame. The specific trial is 24_16_16, (male rat 24, session 16, trial 16).

**Video 3**. Behavior during a single danger trial

Video shows the 75 sequential frames for a danger trial. Frames 1-25 are background and 26-75 are danger cue presentation. Observer judgment is shown in the top right for each frame. The specific trial is 5_16_11, (female rat 5, session 16, trial 11).

**Video 4**. Behavior during a single danger trial

Video shows the 75 sequential frames for a danger trial. Frames 1-25 are background and 26-75 are danger cue presentation. Observer judgment is shown in the top right for each frame. The specific trial is 4_16_3, (male rat 4, session 16, trial 3).

## Materials and Methods

### Subjects

Twenty-four adult Long Evans rats (12 female) weighing 196-298g arrived from Charles River Laboratories on postnatal day 55. Rats were single-housed on a 12-hr light cycle (lights off at 6:00pm) and maintained at their initial body weight with standard laboratory chow (18% Protein Rodent Diet #2018, Harlan Teklad Global Diets, Madison, WI). Water was available *ad libitum* in the home cage. All protocols were approved by the Boston College Animal Care and Use Committee and all experiments were carried out in accordance with the NIH guidelines regarding the care and use of rats for experimental procedures.

### Behavior apparatus

The apparatus for Pavlovian fear discrimination consisted of four individual chambers with aluminum front and back walls, clear acrylic sides and top, and a grid floor. LED strips emitting 940 nm light were affixed to the acrylic top to illuminate the behavioral chamber for frame capture. 940 nm illumination was chosen because rats do not detect light wavelengths exceeding 930 nm <u>(Nikbakht and Diamond, 2021)</u>. Each grid floor bar was electrically connected to an aversive shock generator (Med Associates, St. Albans, VT). An external food cup, and a central port equipped with infrared photocells were present on one wall. Auditory stimuli were generated with an Arduino-based device and presented through two speakers mounted on the ceiling.

### Pellet exposure and nose poke shaping

Rats were food-restricted and specifically fed to maintain their body weight throughout behavioral testing. Each rat was given four grams of experimental pellets in their home cage in order to overcome neophobia. Next, the central port was removed from the experimental chamber, and rats received a 30-minute session in which one pellet was delivered every minute. The central port was returned to the experimental chamber for the remainder of behavioral testing. Each rat was then shaped to nose poke in the central port for experimental pellet delivery using a fixed ratio schedule in which one nose poke into the port yielded one pellet. Shaping sessions lasted 30 min or until approximately 50 nose pokes were completed. Each rat then received 6 sessions during which nose pokes into the port were reinforced on a variable interval schedule. Session 1 used a variable interval 30 s schedule (poking into the port was reinforced every 30 s on average). All remaining sessions used a variable interval 60 s schedule. For the remainder of behavioral testing, nose pokes were reinforced on a variable interval 60 s schedule independent of cue and shock presentation.

### Cue pre-exposure

Each rat was pre-exposed to the three cues to be used in Pavlovian discrimination in one session. Auditory cues consisted of repeating motifs of broadband click, phaser, or trumpet. This 37 min session consisted of four presentations of each cue (12 total presentations) with a mean inter-trial interval (ITI) of 2.5 min. Trial type order was randomly determined by the behavioral program and differed for each rat, each session.

### Pavlovian fear discrimination

Each rat received sixteen, 48-minute sessions of fear discrimination. Each session consisted of 16 trials, with a mean ITI of 2.5 min. Auditory cues were 10 s in duration. Each cue was associated with a unique foot shock probability (0.5 mA, 0.5 s): danger, *p*=1.00; uncertainty, *p*=0.25; and safety, *p*=0.00. Foot shock was administered 2 s following the termination of the auditory cue on danger and uncertainty-shock trials. Auditory identity was counterbalanced across rats. Each session consisted of four danger trials, two uncertainty-shock trials, six uncertainty-omission trials, and four safety trials. Trial type order was randomly determined by the behavioral program and differed for each rat, each session.

### Calculating suppression ratio

Time stamps for cue presentations, shock delivery, and nose pokes (photobeam break) were automatically recorded by the Med Associates program. Baseline nose poke rate was calculated for each trial by counting the number of pokes during the 20-s pre-cue period and multiplying by 3. Cue nose poke rate was calculated for each trial by counting the number of pokes during the 10-s cue period and multiplying by 6. Nose poke suppression was calculated as a ratio: (baseline poke rate – cue poke rate) / (baseline poke rate + cue poke rate). A suppression ratio of ‘1’ indicated complete suppression of nose poking during cue presentation relative to baseline. A suppression ratio of indicated ‘0’ indicates equivalent nose poke rates during baseline and cue presentation. Gradations in suppression ratio between 1 and 0 indicated intermediate levels of nose poke suppression during cue presentation relative to baseline. Negative suppression ratios indicated increased nose poke rates during cue presentation relative to baseline.

### Frame capture system

Behavior frames were captured using Imaging Source monochrome cameras (DMK 37BUX28; USB 3.1, 1/2.9” Sony Pregius IMX287, global shutter, resolution 720×540, trigger in, digital out, C/CS-mount). Frame capture was triggered by the Med Associates behavior program. The 28V Med Associates pulse was converted to a 5V TTL pulse via Adapter (SG-231, Med Associates, St. Albans, VT). The TTL adapter was wired to the camera’s trigger input. Captured frames were saved to a PC (OptiPlex 7470 All-in-One) running IC Capture software (Imaging Source). Frame capture began precisely 5 s before cue onset and continued throughout 10-s cue presentation. Frames were captured at a rate of 5 per second, with a target of capturing 75 frames per trial (5 frames/s x 15s = 75 frames), and 1200 frames per session (75 frames/trial x 16 trials = 1200 frames).

### Post-acquisition frame processing

We aimed to capture 1200 frames per session, and selected sessions 2, 8, and 16 for hand scoring. A Matlab script sorted the 1200 frames into 16 folders, one for each trial, each containing 75 frames. Each 75-frame trial was made into a 75-slide PowerPoint presentation to be used for hand scoring.

### Anonymizing trial information

A total of 1,152 trials of behavior were scored from the 24 rats over the 3 sessions of discrimination (16 trials per session). We anonymized trial information in order to score behavior without bias. The numerical information from each trial (session #, rat # and trial #) was encrypted as a unique number sequence. A unique word was then added to the front of this sequence. The result was that each of the 1,152 trials was converted into a unique word+number sequence. For example, trial ac01_02_07 (rat #1, session #2 and trial #7) would be encrypted as: abundant28515581. The 1,152 trials were randomly assigned to 1 of the 5 observers. The result of trial anonymization was that observers were completely blind to subject, trial type, and session number. Further, random assignment meant that the 16 trials composing a single session were scored by different observers.

### Behavior categories and definitions

Frames were scored as one of ten mutually exclusive behavior categories, defined as follows:

#### Background

Specific behavior cannot be discerned because the rat is turned away from the camera or position of forepaws is not clear, or because the rat is not engaged in any of the other behaviors.

#### Cup

Any part of the nose above the food cup but below the nose port.

#### Freeze

Arched back and stiff, rigid posture in the absence of movement, all four limbs on the floor (often accompanied by hyperventilation and piloerection). Side to side head movements and up and down head movements that do not disturb rigid posture are permitted. Activity such as sniffing or investigation of the bars is not freezing. Freezing, as opposed to pausing, is likely to be 3 or more frames (600+ ms) long.

#### Groom

Any scratching, licking, or washing of the body.

#### Jump

All four limbs off the floor. Includes hanging which is distinguished when hind legs are hanging freely.

#### Locomote

Propelling body across chamber on all four feet, as defined by movement of back feet. Movement of back feet with front feet off the floor is rearing.

#### Port

Any part of the nose in the port. Often standing still in front of the port but sometimes tilting head sideways with the body off to the side of the port.

#### Rear

One or two hind legs on the grid floor with both forepaws *o*ff the grid floor and not on the food cup. Usually (not always) stretching to full extent, forepaws usually (not always) on top of side walls of the chamber, often pawing walls; may be accompanied by sniffing or slow side-to-side movement of head. Does not include grooming movements or eating, even if performed while standing on hind legs.

#### Scale

All four limbs off the floor but at least two limbs on the side of the chamber. Standing on the food cup counts as scaling.

#### Stretch

Body is elongated with the back posture ‘flatter’ than normal. Stretching is often accompanied by immobility, like freezing, but is distinguished by the shape of the back.

### Frame scoring system

Frames were scored using a specific procedure. Frames were first watched in real time in Microsoft PowerPoint by setting the slide duration and transition to 0.19 s, then playing as a slideshow. Behaviors clearly observed were noted. Next, the observer went through the 75 frames scoring one behavior at a time. A standard scoring sequence was used: port, cup, rear, scale, jump, groom, freeze, locomote, and stretch. When the specific behavior was observed in a frame, that frame was labeled. Once all behaviors had been scored, the video was re-watched for freezing. The unlabeled frames were then labeled ‘background’. Finally, all background frames were checked to ensure they did not contain a defined behavior.

### Inter-observer reliability

To assess inter-observer reliability, we selected 12 trials from outside session 2, 8, and 16, six from females and six from males. Each of our five observers scored these 12 trials, interweaving the 12 comparison trials with the primary data trials. As a result, each observer scored 900 comparison frames which were then used to assess inter-observer reliability.

### Statistical analyses

Analysis of variance (ANOVA) was performed for body weight, baseline nose poke rate, suppression ratios, and specific behaviors. Sex was used as a factor for all analyses. Cue, session, and time were used as factors when relevant. Univariate ANOVA following MANOVA used a Bonferroni-corrected p value significance of 0.0055 (0.05/9) to account for the nine quantified behaviors. Multiple analysis of variance (MANOVA) was performed for the nine quantified behaviors with factors of sex, cue, and time. Pearson’s correlation coefficient was used to examine the relationship between baseline nose poke rate and body weight, baseline nose poke rate and cue discrimination, as well as the relationship between danger cue-elicited behaviors during early and late cue presentation in session. Within-subject comparisons were made using 95% bootstrap confidence intervals with the Matlab bootci function. Comparisons were said to differ when the 95% bootstrap confidence interval did not contain zero. Between subject’s comparisons were made using independent samples t-test.

