## Supplemental Figures for "A fear conditioned cue orchestrates a suite of behaviors"

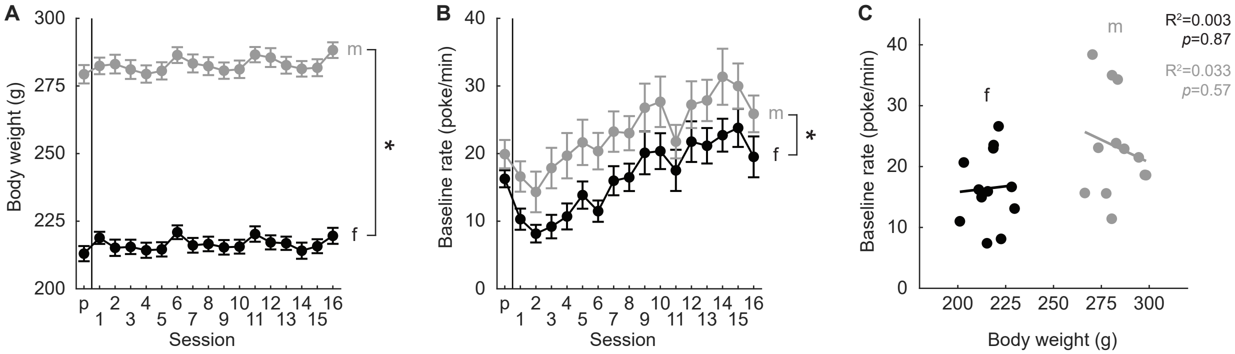


**Figure 1 - Supplemental Figure 1**. Body weight and baseline nose poke rate.

(**A**) Analysis of variance (ANOVA) for body weight (g) [factors: sex and session] revealed significant main effects of session (F(16,352) = 29.58, p = 1.90 x 10-55), sex (F(1,22) = 287.54, p = 4.07 x 10-14), and a significant session x sex interaction (F(16,352) = 2.20, p = 0.005). Mean ± SEM body weights in grams (y-axis) of males (gray) and females (black) from pre-exposure through session 16. (**B**) Baseline nose poke rates (poke/min) decreased during discrimination sessions 1 and 2, then increased over the remaining sessions. Males poked at higher baseline levels across all sessions. Analysis of variance (ANOVA) for baseline nose poke rate (poke/min) [factors: sex and session] revealed significant main effects of session (F(16,352) = 19.30, p = 4.44 x 10-39) and sex (F(1,22) = 5.10, p = 0.034). Mean baseline pose rate (y-axis) of males and females from sessions 1-16. (**C**) Baseline nose poke rate plotted against body weight for all individuals. There was no relationship between the two measures in either female or male rats. *paired samples t-test p<0.05.

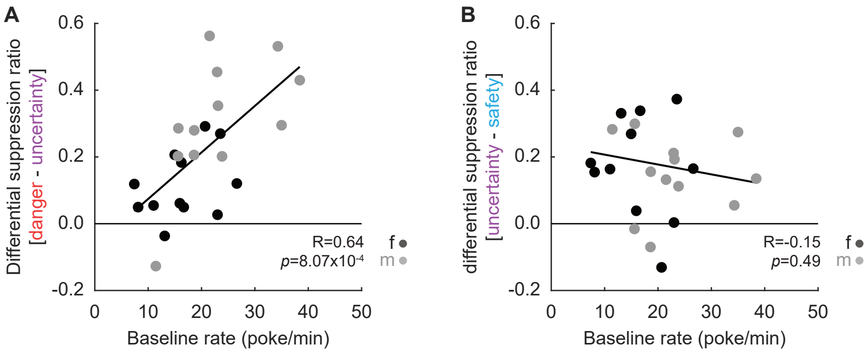


**Figure 1 - Supplemental Figure 2**. Nose poke x discrimination.

Correlations between baseline nose poke rate and differential suppression ratios for **(A)** danger and uncertainty, and **(B)** uncertainty and safety are shown. Individuals represented by black (female) and gray (male) circles. R and *p* values from Pearson's correlation coefficient reported.

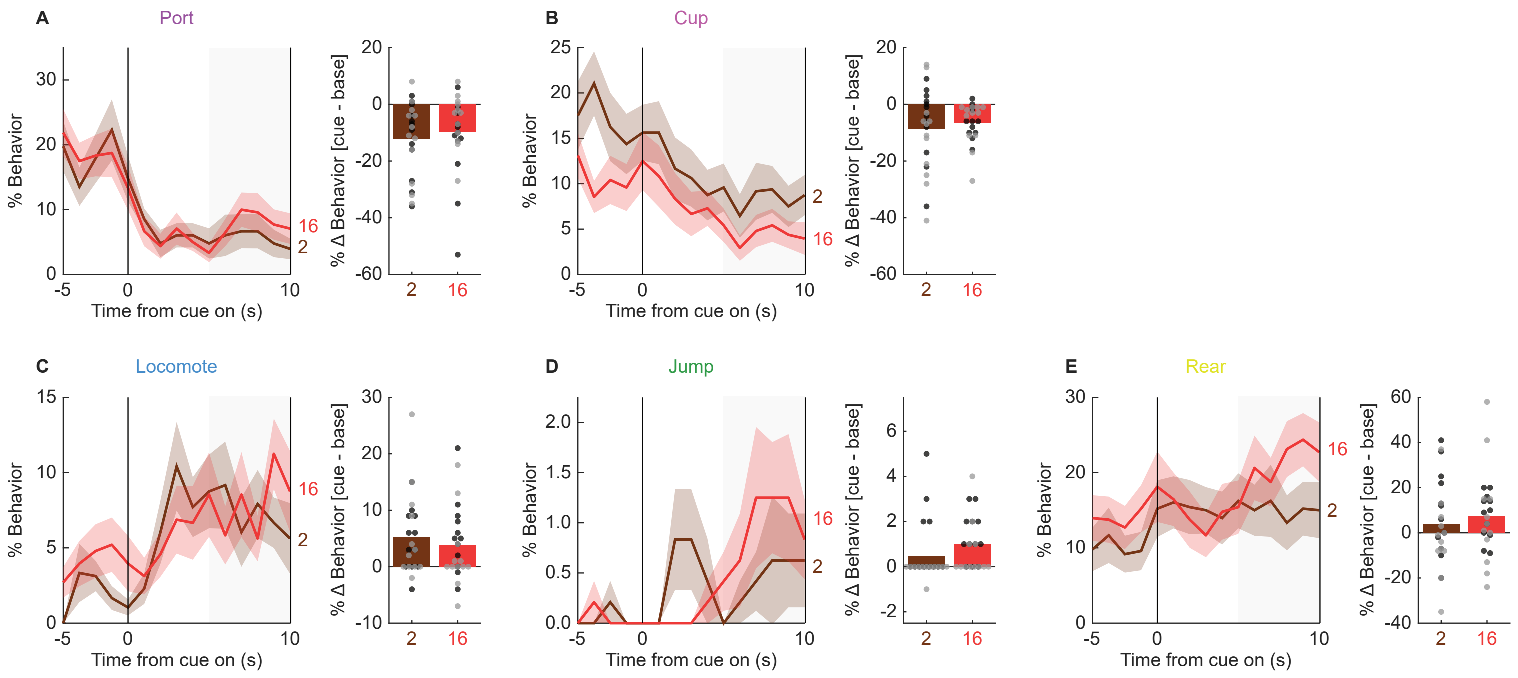


**Figure 5 - Supplemental Figure 1**. Comparions of session 16, danger-specific behaviors during sessions 2 and 16.

Mean ± SEM percent behavior from 5s prior through 10-s danger presentation is shown for **(A)** port, **(B)** cup, **(C)** locomote, **(D)** jump, and **(E)** rear, for session 2 (dark brown) and 16 (red). Adjacent plots show mean % change in behavior from baseline to last 5 s of danger presentation for all rats during session 2 (dark brown) and 16 (red). Individuals data points shown (females, black and males gray).

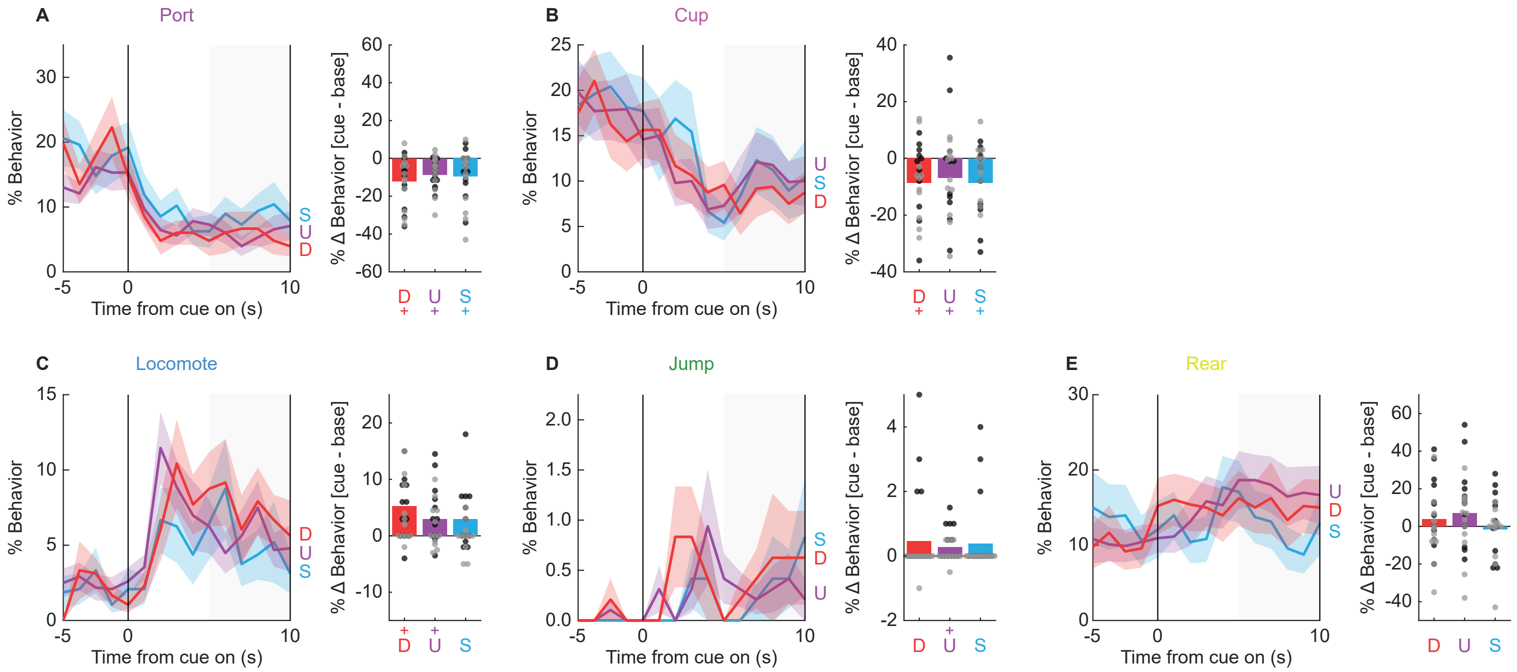


**Figure 5 - Supplemental Figure 2**. Session 16, danger-specific behaviors during session 2.

Mean ± SEM percent behavior from 5s prior through 10-s cue presentation is shown for **(A)** port, **(B)** cup, **(C)** locomote, **(D)** jump, and **(E)** rear, danger (red), uncertainty (purple), and safety (blue). Adjacent plots show mean % change in behavior from baseline to last 5 s of cue presentation for all rats. Individuals data points shown (females, black and males gray). +95% bootstrap confidence interval does not contain zero.

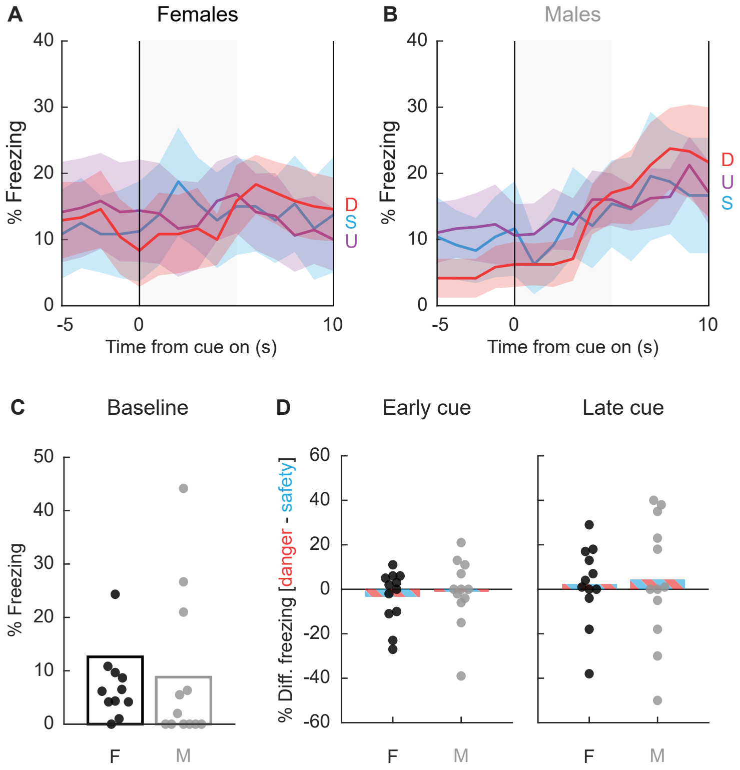


**Figure 6 - Supplemental Figure 1**. Session 2 freezing.

Mean ± SEM percent freezing from 5s prior through 10-s cue presentation is shown for danger (red), uncertainty (purple), and safety (blue) for (**A**) females, and (**B**) males. ANOVA [factors: sex, cue, and interval] found no significant main effect or interaction for cue (Fs < 1.4, *ps* > 0.1). Instead, ANOVA only revealed a significant main effect of interval and a significant sex x interval (Fs > 2.5, *ps* < 0.01). Rats increased freezing to all cues over presentation and this increase was greatest in males. (**C**) Baseline freezing was equivalent in females and males and neither sex showed differential freezing to danger and safety during either early or late cue periods. Mean % differential freezing to danger and safety plotted (**D**). Individuals data points shown (females, black and males gray).
